# Social integration in temporal multiplex association networks predicts offspring survival in female Geoffroy’s spider monkeys (*Ateles geoffroyi*)

**DOI:** 10.1101/2025.10.20.683511

**Authors:** Cristina Jasso-del Toro, Sandra E. Smith Aguilar, Silvia Ruiz Velasco, Xavier St-Amant, Filippo Aureli, Sophie Calmé, Colleen M. Schaffner, Gabriel Ramos-Fernandez

## Abstract

While sociality is considered an important factor influencing female reproductive success, it is unclear how temporal social dynamics relate to it. We address this by evaluating the influence of social integration and its stability over time on offspring production and survival in Geoffroy’s spider monkeys (*Ateles geoffroyi*). Using 52,670 subgroup scan samples collected between 1997 and 2020, we compiled yearly association matrices among the sexually mature individuals of a wild group of spider monkeys, based on spatio-temporal co-occurrence in subgroups. We built Bayesian edge-weight models with the association matrices and generated 1000 samples of each yearly network, which we then used as layers in temporal multiplex networks. For each female with adult tenure ≥ 5 years (n= 17), we estimated strength and eigenvector versatility in the multiplex network that comprised the last 5 years of her tenure. We also estimated six (monolayer) centrality-based metrics in the yearly networks of females with adult-tenure ≥ 1 year and at least one recorded offspring born in the group (n=22). We tested the influence of these eight indicators of social integration as predictors of offspring production and survival. Results showed overall positive relationships between social integration and offspring survival, but not offspring production. The former could be due to better access to food resources or protection from within-group aggression. Our temporal multilayer approach contributed to the body of evidence regarding the fitness benefits of sociality.

**Significance Statement:** Despite the growing evidence of the influence of sociality on female fitness, there is limited information on how temporal variation in social dynamics relates to female reproductive success. We used long-term data on spatio-temporal co-occurrence in subgroups of spider monkeys to estimate network centrality metrics. We found that females with higher and more stable social integration had higher offspring survival, particularly for female offspring who reached age 5 (the onset of sexual maturity). Our study offers insights into the relationship between association and offspring survival in species with a high degree of fission-fusion dynamics and female dispersal.

## Introduction

There is a growing body of evidence from different group-living mammals highlighting the impact of sociality on female reproductive success and survival (Ostner and Schülke 2018). Females who maintain close and stable relationships with favoured partners, or spend more time in close proximity to or grooming others, tend to have (1) longer life spans (chacma baboons, *Papio hamadryas ursinus*, Silk et al. 2010; rhesus macaques, *Macaca mulatta*, Ellis et al. 2019; yellow baboons, *Papio cynocephalus*, Archie et al. 2014, Campos et al. 2020b, Lange et al. 2023; white-faced capuchins, *Cebus capucinus,* Kajokaite et al. 2022), (2) higher offspring survival (yellow baboons, Silk et al. 2003; feral horses, *Equus caballus*, Cameron et al. 2009; bottlenose dolphins, *Tursiops sp*., Frère et al 2010) and (3) higher birth rates (wild house mice, *Mus domesticus*, Weidt et al. 2008; feral horses, Cameron et al. 2009). Some studies also point to negative (Barocas et al. 2011; Ellis et al. 2019; Menz et al. 2020) or more nuanced (Sabol et al. 2020) relationships between social behavior and fitness.

There are many potential mechanisms behind this relationship (reviewed by: Massen et al. 2010; Ostner and Schülke 2018; Thompson 2019; Dunbar 2024). For example, close social relationships with a few preferred partners can decrease stress levels (Wittig et al. 2008), leading to increased survival (Rakotoniaina et al. 2017). Associating with or being tolerant towards multiple partners can provide protection from predators, improve thermoregulation, and reduce injury risk, thereby enhancing survival (Kern and Radford 2016; Campbell et al. 2018; Pavez-Fox et al. 2022). Similarly, individuals with multiple and strong social relationships may share information about food locations, enabling access to resources (Carter et al. 2016), a crucial factor for female reproductive success and survival (Gittleman and Thompson 1988; Ostner and Schülke 2018). Information sharing is particularly relevant for species with a high degree of fission-fusion dynamics where group members frequently split into and join subgroups of different size and composition (Aureli et al. 2008) as näive individuals may be exposed to information on foraging sites discovered by group members in other subgroups upon meeting, for example at sleeping sites (e.g. Aureli et al. 2008; Palacios-Romo et al. 2019; Papageorgiou et al. 2024). Females can also increase their reproductive success by forming coalitions with others for defence against harassment from other group members and infanticidal males, which can impact food intake and risk of injury or death of either females or their offspring (e.g., Sterck et al. 1977; Palombit et al. 1997; Cameron et al. 2009).

Researchers using social network analyses found centrality metrics predicting female fitness in multiple mammalian species (wild rock hyraxes, *Procavia capensis*, Barocas et al. 2011; Barbary macaques, *Macaca sylvanus*, Lehmann et al. 2015; chacma baboons, Cheney et al. 2016; yellow-bellied marmots, *Marmota flaviventer*, Montero et al. 2021; spotted hyenas, *Crocuta crocuta*, Turner et al. 2021). Here we use social network analysis to examine female “social integration”, defined as how well an individual is embedded in the network of affiliative relationships of its social group (in our case proxied by association in the same subgroup). Social integration, commonly quantified by using centrality metrics, captures direct (strength centrality, degree and number of weak connections) and indirect connectedness (betweenness, closeness and eigenvector centrality; Ellis et al. 2019; Snyder-Mackler et al. 2020; Soben et al. 2023).

Social integration is prone to vary over time due to changes in individual attributes (e.g., age class, group tenure) as well as social, demographic, and ecological events (e.g., deaths, births, migration, environmental variation; Williams et al. 2017; Shizuka and Johnson 2019; Evans et al. 2020b; Rathke and Fischer 2021; Xia et al. 2021), with potential consequences to individual fitness (e.g., Rathke and Fischer 2021; Siracusa et al. 2023). Changes in group composition due to the loss of individuals (e.g., death, emigration) may lead to the establishment of new connections, which can vary in strength depending on the social integration of each individual prior to the change, or may result in a reduced number of associates (Shizuka and Johnson 2019; Evans et al. 2020a). Similarly, immigration involves new potential interactions for immigrants and residents, impacting individual (and possibly group-wise) patterns of social integration (e.g., Xia et al. 2021). Environmental factors such as habitat disturbances (e.g., fires and hurricanes; Lantz and Karubian 2017; Testard et al. 2021), resource availability (Henzi et al. 2009; Lantz and Karubian 2017; Testard et al. 2021), and temperature (Rat et al. 2020; Rothberg et al. 2024) can also change social connectivity between individuals in species-specific ways. For example, it increases proximity and decreases aggression toward conspecifics after a hurricane (Testard et al. 2021).

While analyses of the relationship between network-based social integration and female fitness largely use monolayer networks (e.g., Barocas et al. 2011; Lehmann et al. 2015; Cheney et al. 2016, but see Sharma et al. 2022), time-aggregated or temporal multiplex networks can potentially reveal complementary aspects of social dynamics (e.g., temporal changes in individual roles in a group; e.g., Vilette et al. 2025). These dynamics can be relevant for individual fitness but may be missed if compressed into a monolayer network (e.g., Bonnell et al. 2020). Thus, we chose a multilayer approach to examine the temporal dynamics of an individual’s network connectivity using a multiplex network to shed light on the link between social integration and fitness. Multiplex networks are multilayer networks where layers can represent different types of social interactions or different time windows of the same interaction type among a given set of individuals (Silk et al. 2018; Finn et al. 2019; De Domenico 2022). Layers are connected to each other through interlayer edges that link individuals to themselves across different interaction types or time windows.

Our study focuses on Geoffroy’s spider monkeys (*Ateles geoffroyi*), a species living in multi-female, multi-male groups with a high degree of fission-fusion dynamics where group members form subgroups that frequently change size and composition throughout the day (Aureli and Schaffner 2008). The distribution of spider monkey social interactions is largely driven by partner sex-age classes (Aureli and Schaffner 2008; Ramos-Fernández et al. 2009; Hartwell et al. 2014) and maternal kinship (Jasso-del Toro et al. 2024). In the spider monkey social system, where contest and scramble competition for food are regulated by fission-fusion dynamics (Asensio et al. 2008, 2009), female association patterns reflect social tolerance and passive attraction to resources of common interest such as food and shelter (Ramos-Fernández et al. 2006; Slater et al. 2009; Smith-Aguilar et al. 2016). Relationship quality and the presence of individuals with novel information seem to influence individual decisions to join or depart from subgroups (Ramos-Fernández and Morales 2014; Busia et al. 2017; Palacios-Romo et al. 2019). Furthermore, female sociality is related to group tenure and reproductive status (Asensio et al. 2015; Shimooka 2015; Riveros et al. 2017). In particular, females associate more with other females, forming weaker relationships among themselves than males do with other males, and may display active avoidance of males (Ramos-Fernández et al. 2009; Slater et al. 2009; Smith-Aguilar et al. 2016). Before sexual maturity, females basically share the social connections of their mothers (Jasso-del Toro et al. 2024). Once they approach sexual maturity (at around 5 years of age) prior to leaving their natal group and right after immigrating into a new group, females tend to have few and weak connections and are subject to aggression from other group-members, particularly other females (Asensio et al. 2008; Shimooka et al. 2008; Riveros et al. 2017). This pattern typically results in low social integration for newly immigrated females, as indicated by low centrality scores (Ramos-Fernández et al. 2009; Smith-Aguilar et al. 2019).

It is possible that the challenges for offspring survival prior to sexual maturity are not equivalent for the two sexes (e.g., Álvarez et al. 2015), and maternal investment may be biased towards one sex as it is the case for male offspring in some primate species with male philopatry like chimpanzees (*Pan troglodytes*; Bădescu et al. 2022) and other spider monkey species (e.g., *A. paniscus*; Symington 1987). Moreover, maternal investment may be modulated by social integration (Soben et al. 2023) with older, more central *A. geoffroyi* females possibly investing more in male than female offspring because male offspring provide higher fitness returns (Trivers and Willard 1973). Consequently, having highly central mothers could confer uneven survival benefits for males and females, either by having differential access to the survival benefits of direct maternal care or of reduced predation or risk of infanticide through vigilance or dilution effects.

The aim of our study was to evaluate the influence of social integration and its stability over time on female reproductive success using temporal multiplex association networks. Although association in the same subgroup does not capture the multidimensionality of social relationships, it can provide an approximation (Whitehead and Dufault 1999; Smith-Aguilar et al. 2019) allowing us to examine temporal dynamics over two decades. We hypothesised that females who are better integrated over time have greater reproductive success. Thus, we predicted that females with higher centrality and females with more stable central positions would tend to have more offspring and higher offspring survival rates. We further predicted that highly central females would have higher male than female offspring survival.

## Methods

### Study site

The study site is located in the *Otoch Ma’ax yetel Kooh* protected area (5,367 ha), near the village of Punta Laguna in the Yucatan peninsula (Mexico; 20°38′ N, 87°38′ W). The site comprises areas of tropical semi-evergreen forest in different successional stages (Ramos-Fernández et al. 2018). The area is typically characterised by a dry season from November to April and a wet season from May to October (Spaan et al. 2021).

### Data collection and study subjects

From January 1997 to December 2020, experienced observers followed monkey subgroups for 4 – 8 hours daily (Ramos-Fernández et al. 2018). Subgroups were defined using a chain rule (Croft et al. 2008), by which individuals within 30 m of a subgroup member were considered to belong to the focal subgroup (Ramos-Fernández 2005). Once a subgroup was found, observers conducted instantaneous scan samples (Altmann 1974) every 20 minutes recording the identity of all subgroup members. The observers recognized each individual by distinctive marks and coloration in their face or body. Overall, the dataset comprised 52,670 scan samples collected over 3,511 observation days over 24 years with a mean of 2,195 (±892) scan samples per year. Because our study involved instantaneous scan samples in the field, it was not possible to record data blind.

A total of 93 individuals were included in the dataset (63 females and 30 males). Birth dates were known (or estimated) for 79 individuals born after 1997. For individuals born prior to 1997, we estimated their minimum age based on size, behavior and, in the case of fully grown females, the presence/absence of infants and juveniles travelling with them. We assumed that females who immigrated into the study group were 6 years old when no offspring travelled with them (i.e., the average age at which natal females emigrated out of the study group); otherwise, we assigned them a minimum age by assuming they were 8 years old at the birth of their oldest accompanying young (this is the age at which the only natal female that did not emigrate had her first offspring). An individual was considered dead if their body was found or if it was seriously injured the last time it was seen. When an individual went unseen for a month but death was not confirmed, we set the last day of observation as the date of disappearance.

### Reproductive success

We evaluated reproductive success for females older than 6 years that were observed in the group for at least one year and had at least one offspring while being part of the group. Twenty-two females fulfilled these criteria, with tenure time (TT; defined as the period between the first and last record of a given female in a scan sample during the study period) ranging from 1.6 to 24 years (Fig. S1). Six of the females were group residents prior to 1997, one was born in the group (i.e., she did not emigrate), and 15 immigrated into the group during the study period. By the end of the study, 11 of the 22 females had disappeared. Data on birth and offspring survival considered for analyses are confined to the 1997-2020 period, not to the full reproductive life of the females. The number of recorded offspring per female ranged from 1 to 10, totalling 77 offspring (38 female, 31 male and 8 of unknown sex).

We used records of births, deaths and disappearances to estimate four indicators of female reproductive success based on the number of known offspring (offspring production) and their survival to 1, 3 and 5 years (offspring survival). We also calculated survival to 5 years for male and female offspring separately. To correct for differences in the period each female could possibly reproduce, we estimated the reproductive years (RY) in the group for each female during the study (see Table 1 for details). RY compensates for the difference between females who already had offspring when data collection began and females who had offspring several months or years after they immigrated (i.e., whose sexual maturity was uncertain before their first pregnancy in the study group).

**Table 1.** Variables in the statistical models to assess the reproductive success of each female.

| Variable | Definition | Description |
| --- | --- | --- |
| **Offspring production**<br>(response variable: discrete) | Number of female/male/total offspring produced | Number of female/male/total offspring born in the group during the study period (01/January/1997 - 31/December/2020). |
| **Offspring survival**<br>(response variable: discrete) | Number of total offspring reaching 1 year | Number of offspring born in the group after 01/January/1997 and before 31/December/2019, that reached 1 year of age. |
|  | Number of total offspring reaching 3 years | Number of offspring born in the group after 01/January/1997 and before 31/December/2017, that reached 3 years of age. |
|  | Number of female/male/total offspring reaching 5 years | Number of female/male/total offspring born in the group after 01/January/1997 and before 31/December/2015, that reached 5 years of age. |
| **Reproductive years**<br>(offset variable) | Number of reproductive years (RY) | RY=RT except for females who immigrated into the group |
| | | during the study period, when<br>$RY = TT - TFP$, where TFP is the<br>period between immigration<br>date and first known pregnancy,<br>estimated 7 months prior to the<br>birth of the first offspring after<br>immigration. |

### Association networks

We constructed networks considering all individuals older than 5 years (the age at which they can be found in subgroups different from those of their mother) in any given year. We defined an association between two individuals as their co-occurrence in the same subgroup in a given instantaneous scan sample. We built Bayesian edge-weight models for each yearly network following the BiSoN framework proposed by Hart et al. (2023) for binary group-based (gambit of the group) data (details in SI). Overall, we built 24 edge-weight models, one for each study year, from which we obtained 1000 association network samples per year (24,000 networks in total). In these network samples, nodes were the individuals present in the study group in a given year and weighted non-directional edges represented association values for each dyad, drawn from the probability distribution for each edge in the model.

### Multiplex temporal networks

We used the yearly association networks as layers to build temporal multiplex networks (multiplexes, hereafter) where each layer represents associations during a year. This was done using normalised association matrices taken from the 1000 samples of each yearly association network produced with the edge-weight models. Normalisation was done by dividing the edge weight of each dyad by the maximum edge weight for that layer (yearly network). In multiplex networks, nodes need not appear in all layers but a node is connected to itself in different layers resulting in a multilayer network structure with intra-layer and inter-layer edges (De Domenico et al. 2013; Kivelä et al. 2014; Finn et al. 2019; Finn, 2021). Given the temporal nature of our multiplexes, inter-layer edges were directed (with weight set to 1), connecting a node to itself in the layer for the following year.

Because metrics in multiplexes can be influenced by the number of layers in the network, we based any multiplex-derived variables on multiplexes with the same number of layers (i.e., the same number of years) for every female. To maximise the number of layers, whilst including as many females as possible, we chose 17 females with group tenure of at least 5 years for the multiplex analysis. This is less than the mean (±SD) tenure of 10.3 years (±7.4; median=8.6) for the 22 selected females, thus allowing us to capture potential variation for females with above- and below-average tenure by including females with TT ranging 5-24 years. For each of the 17 females, we used the final 5 layers (years) of their group tenure to build their corresponding multiplex. By choosing the final years of tenure (rather than initial or middle) we aimed to capture the period when each female had had the most time to integrate (there is no evidence for a non-reproductive period at the end of tenure for any of the studied females). Because we had 1000 samples from each yearly network, we built 1000 5-layer multiplexes for each female. This way, we obtained one thousand samples for each multiplex metric derived from the corresponding 5-layer multiplexes for each female. Multiplexes were built and analysed using the MuxViz 3.1 (De Domenico et al. 2015a; De Domenico 2022) package in R (v. 4.2.0).

### Social integration

We operationalized social integration by means of eight centrality-based metrics (Table 2). Four of these pertained to a female’s centrality in each of the yearly networks or versatility in the multiplex networks relative to others, to capture how well she is connected within her social network. Two metrics (CV strength and CV eigenvector centrality) capture stability in terms of the consistency in centrality scores through time, and two metrics (Δ strength and Δ eigenvector centrality) represent change in centrality scores between early and late tenure for each female. Six of the eight metrics were based on calculating monolayer strength and eigenvector centrality in the yearly association networks for the 22 females across their tenure, while the remaining two metrics were multiplex strength and eigenvector versatility calculated for 17 females in their 5-layer multiplexes. Since we had 1000 samples of each yearly and multiplex network, we obtained 1000 samples from each of the eight social integration metrics for each female.

**Table 2.**
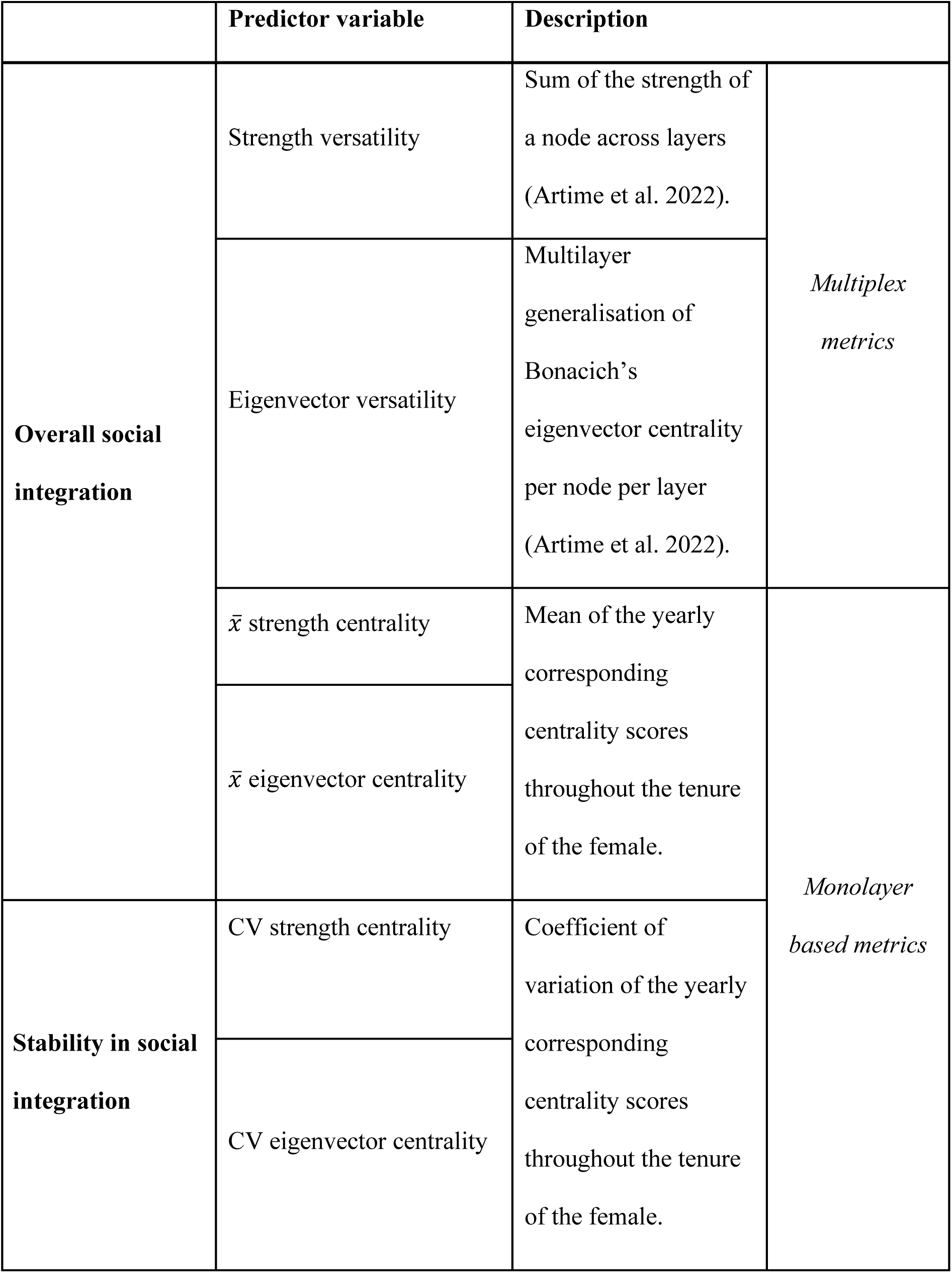
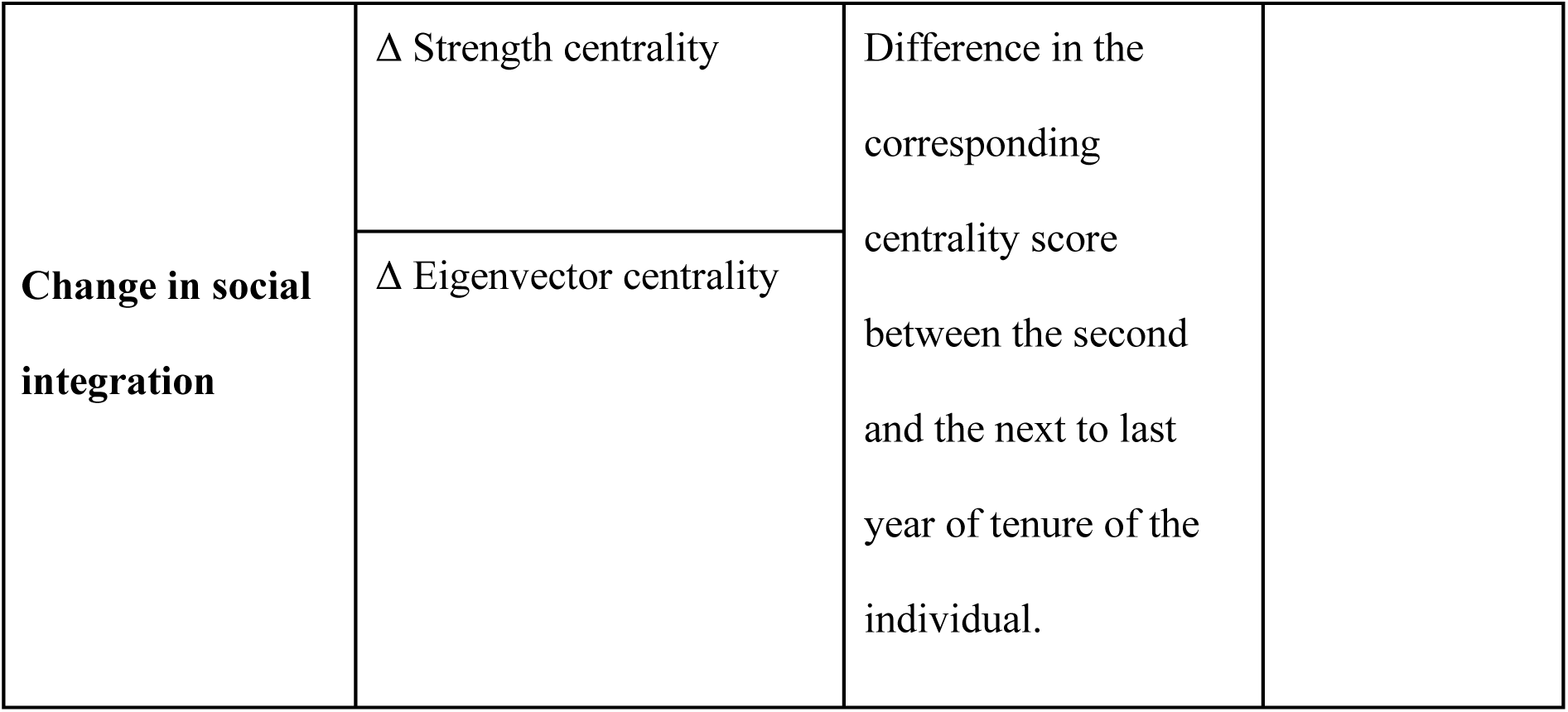
Metrics of social integration used as predictor variables in the statistical models.

We focused on strength and eigenvector centrality because they represent direct and indirect connection patterns, with potentially different consequences for individuals (Brent 2015). Strength highlights nodes with relatively numerous and/or strong connections, accounting for the sum of the weights of all their edges, whereas eigenvector centrality extends to the indirect connections of a node, by identifying the leading eigenvector of the adjacency matrix, giving high scores to nodes with many edges or connected to highly connected others (nodes which themselves have high eigenvector centrality; Newman 2004; Hasenjager and Dugatkin 2015). Strength and eigenvector versatility are mathematical extensions for multilayer networks of the equivalent centrality measures, which account for interlayer edges, producing values calculated from the position of the node in the whole multiplex (De Domenico et al. 2015b).

The range of strength centrality values in any given network is influenced by the number of nodes, edges and by the edge weights. Hence, comparison of absolute centrality values between different networks does not necessarily reflect change in the patterns of social integration which are based on the relative connectedness of an individual with respect to others in the same network. Consequently, for each yearly or multiplex network sample, we normalised the centrality or versatility values using min-max scaling. We then used the normalised values from each of the 1000 network samples, as individual centrality or versatility scores for all our analyses instead of the absolute values (Table 2).

For each series of yearly networks over the TT of a female, we calculated the female’s mean score for each of the two centralities and used them as indicators of overall social integration (*x̅* strength centrality and *x̅* eigenvector centrality; Table 2), hence obtaining 1000 mean values for each centrality metric per female. Similarly, we calculated strength and eigenvector versatility for each female on her corresponding multiplex samples obtaining 1000 values of each versatility per female. Regarding stability in social integration, we calculated the coefficient of variation of both centrality metrics (CV strength centrality and CV eigenvector centrality) across the yearly networks for each female for each of her 1000 series, obtaining 1000 coefficients of variation per centrality metric per female. We also estimated change in social integration over time as the difference in centrality scores between the second and the next to last year of tenure time for each female (Δ strength centrality and Δ eigenvector centrality). Positive values indicate a higher centrality score by the end of the tenure time than at the beginning and negative values indicate the opposite. We used the second and next-to-last years of tenure to consider change in centrality scores based on 12 months of data (as a female’s centrality in the first and last yearly network of her tenure time can be derived from a variable number of months of data depending on the date when she entered/ceased to be in the group). Consequently, Δ centralities were only available for females with residency time > 3 years (n=18). Change in social integration was also calculated for each of the 1000 series of centrality scores mentioned before, resulting in 1000 Δ centrality values per female.

### Statistical analyses

To test the influence of social integration on reproductive success, we mostly fit generalised linear models (GLMs) with 1 response and 1 predictor variable, where response variables were the indicators of offspring production and offspring survival (Table 1), while predictor variables were the indicators of overall social integration and its stability (Table 2). We did not include all predictors in one model because of high correlations among them.

We only used GLMs with >1 predictor variable for change in social integration, to account for potential effects derived from the portion of the group tenure represented. Because Δ strength and Δ eigenvector centrality are based on the whole tenure period for some but not all females, it was possible that those who were already residents by the beginning of the study (and therefore already integrated) would undergo less change in centralities than those who began their integration process during the study. The GLMs included the predictors Δ strength centrality or Δ eigenvector centrality together with the predictor “residency status”, and the interaction between both predictors. We classified females as “immigrant” if they entered the group during the study period (n=13) or “resident” if already in the group at the onset of the study (n=5).

In every GLM, we used a quasi-poisson distribution to account for underdispersion and included the number of reproductive years for each female (as described in Table 1) as an offset variable to correct for differences between reproductive periods for each female. GLMs were run with the “glm” function from the “stats” package in R (R Core Team, 2023). Each GLM was run 1000 times using the 1000 samples for each social integration variable, so that we obtained a distribution for each of the model estimates. Estimates from the 1000 repeated models and the uncertainty around them were combined using the *pool* function in the R package *mice* (van Buuren and Groothuis-Oudshoorn 2011) to report a single estimate for each model.

## Results

We found significant relationships between social integration and offspring survival, but not with offspring production (Fig. 1, Table S2). Females with higher strength and eigenvector versatilities had more offspring that survived to 1 (Fig. 2a), 3 (Fig. 2b) and 5 years (Fig. 2c), especially female offspring that survived to 5 years (Fig. 2d). An increase of one standard deviation in strength (σ= 0.18) and eigenvector versatility scores (σ=0.26) is associated with an increase of a) 11% and 13%, respectively, in the number of offspring surviving to 1 year, b) 21% and 18% in the number of offspring surviving to 3 years, c) 51% and 42% in the number of offspring surviving to 5 years, and d) 84% and 70% in the number of female offspring surviving to 5 years.

**Fig. 1.**
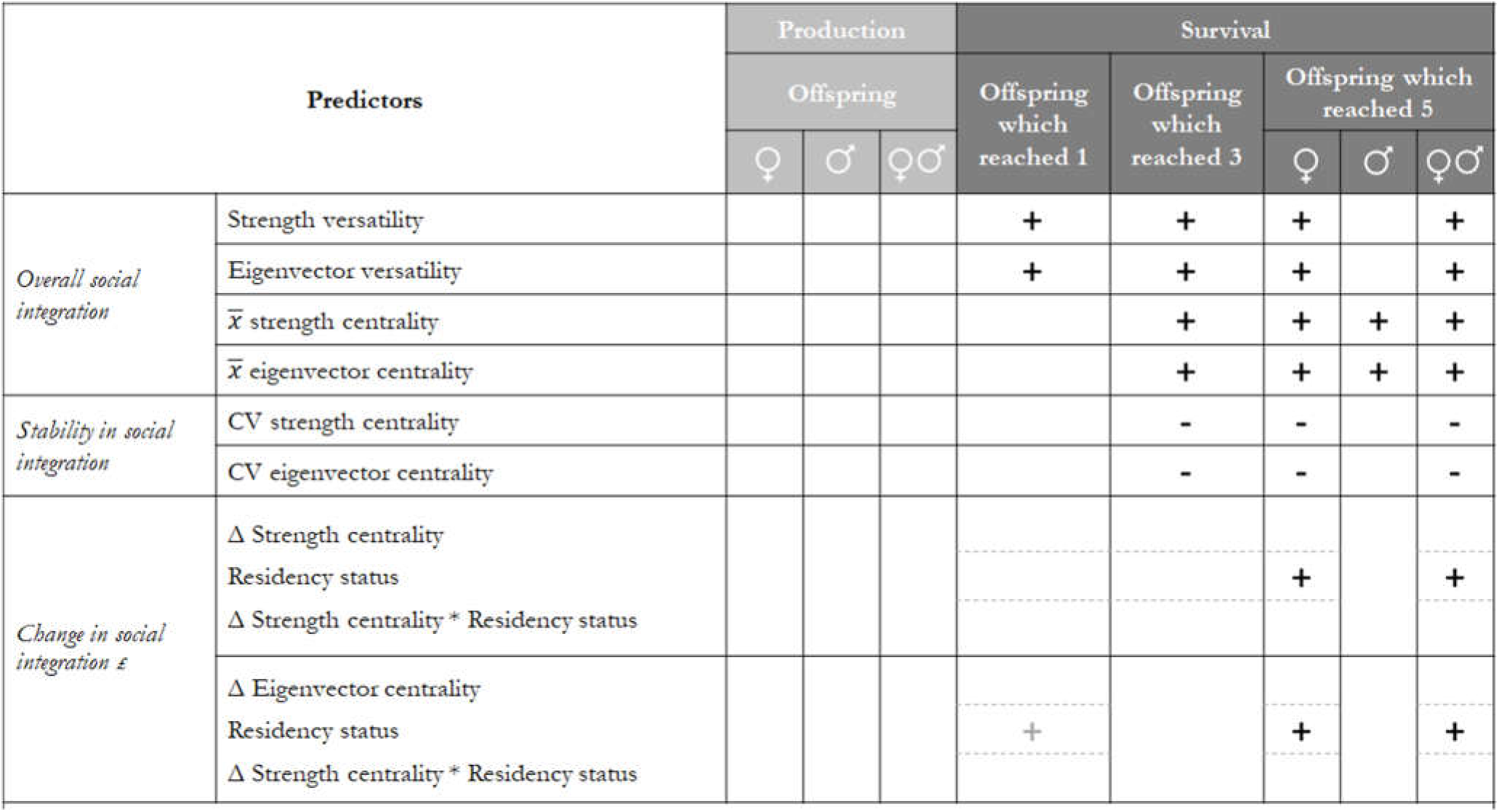
Model matrix indicating significant (black; p<0.05) or marginally significant (grey; p=0.06) positive (+) or negative (-) relationships between social integration (predictors; rows) and reproductive success (columns). Each cell represents a given model (empty cells denoting no relation). *£* Models with two predictors and their interaction. Here, symbols indicate associations for each predictor in the model. See Tables S2 and S3 for details on all models.

**Fig. 2.**
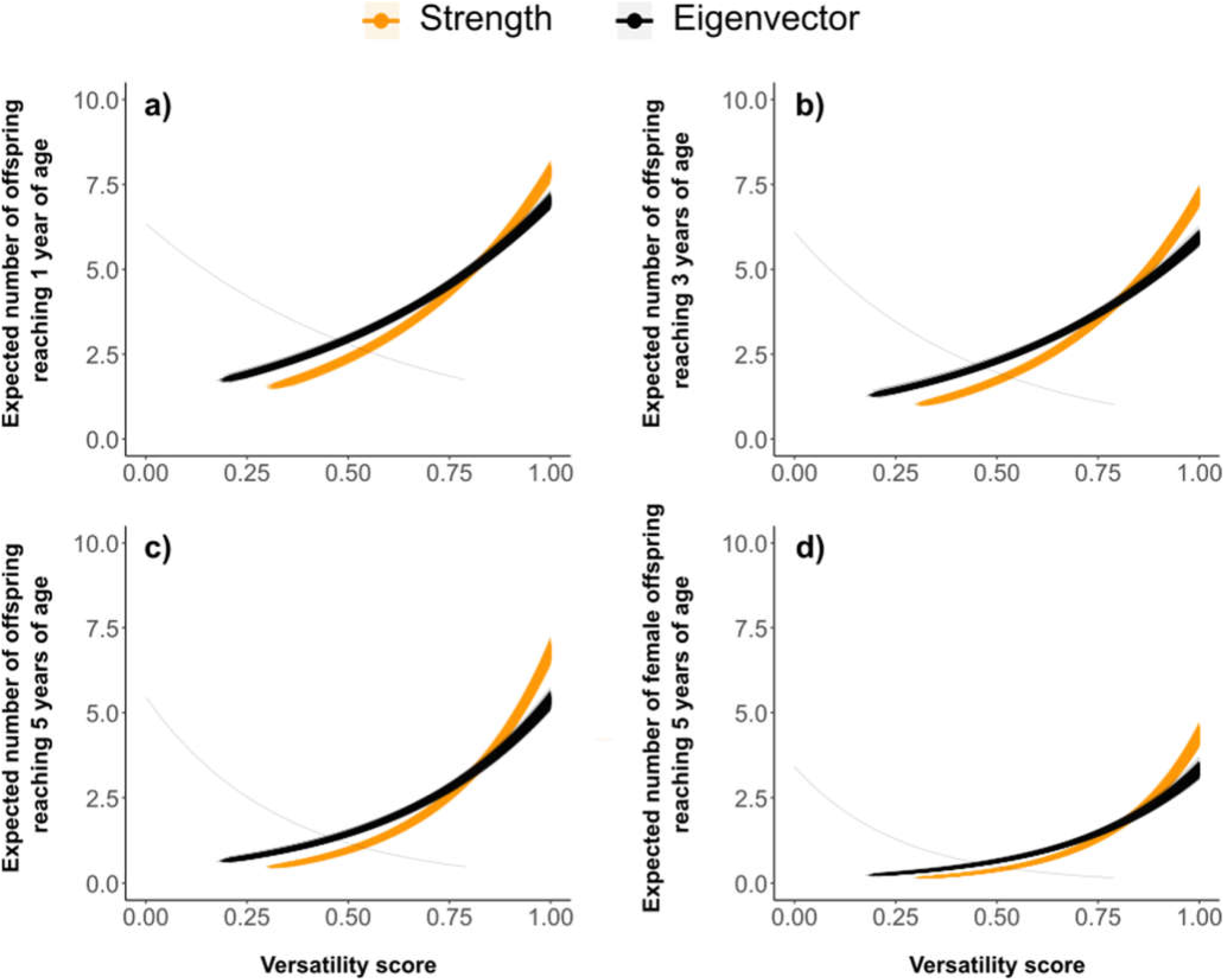
GLM quasi-poisson trend lines for strength (orange) and eigenvector versatility (black and gray) showing their relationships with the expected number of offspring to survive to 1 year (a), 3 years (b), and 5 years (c) and the expected number of female offspring who survived to 5 years. Each trend line represents one of the 1,000 GLM models. For eigenvector versatility only 1 (gray line) out of 1,000 models exhibited a negative trend

Mean centralities rendered similar results: females with higher strength and eigenvector centrality scores had more offspring that survived to 3 (Fig. S2a) and 5 years (Fig. S2b) and more female and male offspring that survived to age 5 (Fig. S2c and S2d). An increase of one standard deviation (σ=0.25) in mean strength and eigenvector centrality scores results in an increase of a) 24% and 23 %, respectively in the number of offspring surviving to 3 years, b) 69% and 69% in the number of offspring surviving to 5 years, c) 99% and 96% in the number of female offspring surviving to 5 years, and d) 44% and 46 % in the number of male offspring surviving to 5 years.

To explore if differences in the number of females included in models using versatilities (n=17) vs. mean centralities (n=22) explained the contrasting results for offspring survival to 1 year and male offspring to 5 years, we ran the monoplex-based models for these variables with mean strength and mean eigenvector centrality as predictors, considering only the 17 females who were included in the multiplex networks. We obtained the same results as for the versatility predictors (i.e. significant relationships for both centralities with offspring survival to 1 year and no significant relationships between these predictors with male offspring survival to 5 years). Therefore, the multiplex-based and monoplex-based social integration models produce the same results if they only include females with TT > 5 years (n=17).

Regarding stability in social integration, we found negative relationships between the coefficient of variation (CV) of strength and eigenvector centralities with total offspring survival to 3 and 5 years, and with female offspring survival to 5 years (Fig. 1, Table S2). Females with less variation in strength and eigenvector centralities over time had more offspring that survived to 3 (Fig. 3a) and 5 years of age (Fig. 3b), and more female offspring that survived to age 5 (Fig. 3c). A decrease of one standard deviation in CV strength (σ= 45) and CV eigenvector centrality scores (σ= 48) (i.e. more stability in social integration) is associated with a) a 25% and 20 % increase, respectively, in the number of offspring surviving to 3 years, b) a 44% and 39% increase in the number of offspring surviving to 5 years, and c) a 63% and 59 % increase in the number of female offspring surviving to 5 years. Even though females could have low social integration scores which are stable over time, it is unlikely that these females are driving the CV results given the negative correlations between CV and mean corresponding centrality metrics (CV strength and *x̅* strength centrality: *r* =-0.77, *p* <0.001; CV eigenvector and *x̅* eigenvector centrality: *r* =-0.79, *p* < 0.001).

**Fig. 3.**
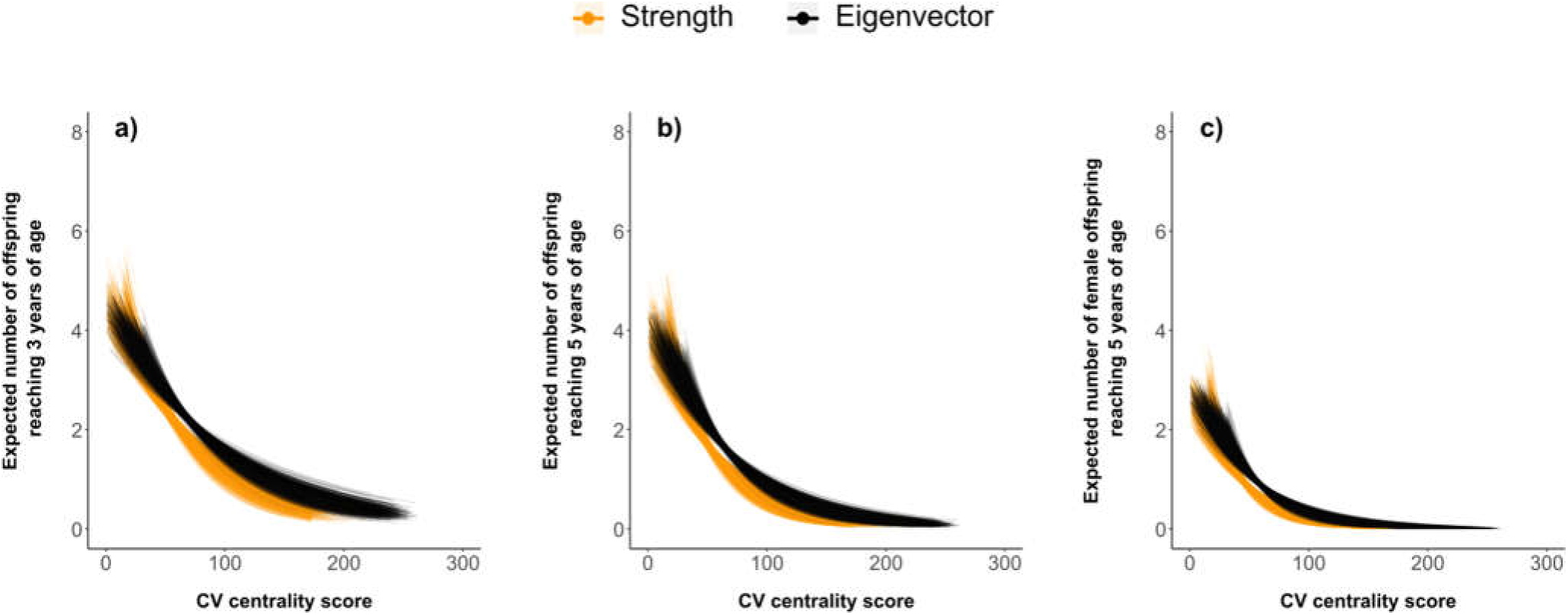
GLM quasi-poisson trend lines (curves) for CV strength (orange) and eigenvector centrality (black) showing their relationship with the expected number of offspring to survive to 3 years (a), 5 years (b) and the expected number of female offspring to survive to 5 years (c). Each trend line represents one of the 1,000 GLM models

Indicators of change in social integration were not significant predictors of either offspring production or offspring survival (Table S3). Although we did not find significant relationships of Δ strength centrality or Δ eigenvector centrality with offspring production or survival, there were significant positive effects of residency status on offspring survival to 5 years and female offspring survival to 5 years. “Resident” females had more offspring and more female offspring that survived to 5 years than females who immigrated into the group during the study period. Residency status was also marginally associated with offspring survival to 1 year in the model including Δ eigenvector centrality (Table S2).

## Discussion

We hypothesised that females who are better socially integrated over time have greater reproductive success. As predicted, females with more central positions and those with more stable positions during group tenure had higher offspring survival, particularly to 5 years, the onset of sexual maturity (but note that 4/20 and 4/10 of the models for overall social integration and stability in social integration, respectively, did not produce significant results). We could not test the interaction effect of overall social integration and stability in social integration on offspring survival because the metrics for both predictors were negatively correlated. However, females with higher social integration tended to have higher stability in their social integration than females with low social integration. Thus, it is more likely that females with higher offspring survival have higher and more stable centrality scores. We did not find support for our prediction that highly central females would have higher male than female offspring survival. On the contrary, we found relationships of social integration and stability in social integration with offspring survival for female offspring using the multiplex and monolayer metrics, whereas for male offspring it was only found with the metrics calculated from the yearly monolayer networks. In contrast with our predictions, change in social integration had no relationship with offspring survival and we did not find significant relationships between any of our social integration metrics and offspring production.

Based on 24 years of data on a wild group of spider monkeys, our study indicates that female social integration, measured with direct and indirect measures of centrality, had a positive relationship with offspring survival. Being highly connected in a spider monkey group can potentially increase access to food resources, which could be particularly important for females to cover the energetic costs associated with gestation and maternal care, as documented for other primate species (Altmann and Samuels 1992; Dufour and Sauther, 2002; Thompson et al. 2012). Spider monkey females carry their youngest offspring for about 19 months, when nursing rejection begins, representing slow offspring development compared to other similarly sized primates (Arbaiza-Bayona et al. 2022). This may be related to a relatively large brain (Street et al. 2017) that continues developing after birth (Leigh 2004) and to high cognitive demands that require high caloric inputs (Heldstab et al. 2022).

Given their high degree of fission-fusion dynamics, spider monkeys can regulate food competition by splitting into smaller subgroups when food is scarce (Asensio et al. 2008; Smith-Aguilar et al. 2016). Despite their daily splitting in multiple subgroups, individuals still interact regularly with many other group members due to frequent changes in subgroup composition, as reflected by highly connected association networks with little evidence of modular substructuring (e.g., Smith-Aguilar et al. 2019). Highly connected networks allow for access to social information (Danchin et al. 2004), which may be particularly relevant given the ephemeral, clumped and heterogeneously dispersed nature of their food sources (Kohles et al. 2022). For instance, Palacios-Romo et al. (2019) found that spider monkeys follow central individuals to guide their foraging decisions. The low rates of within group aggression in spider monkeys (Slater et al. 2009), suggests they may be particularly tolerant to the presence of group members and thereby may access more social information (Duboscq et al. 2016). Furthermore, food abundance can drive association among females by being attracted to the same food patches (Ramos-Fernández et al. 2009; Smith-Aguilar et al. 2016). The relationship of female high social integration and more stable social integration with higher offspring survival could be mediated by central females having better access to information about food sources allowing for enough food intake to cover the energetic costs of lactation, offspring carrying and overall offspring care.

Being a more central individual may also facilitate protection from predation or social support when subject to attacks. Although uncommon, predation (Busia et al. 2018) and lethal intra-group aggression (Campbell 2006; Valero et al. 2006), including infanticide (Gibson et al. 2008; Álvarez et al. 2015), do occur in spider monkeys (Aureli and Schaffner 2008). High centrality in our study mostly indicates having stronger connections as a result of spending time together with group members more frequently. Although strength and eigenvector centralities/versatilities are also influenced by the number of edges of a node, the yearly time-scale of our analyses results in networks where most individuals are connected to all others. Consequently, variation in centrality/versatility scores is mostly due to differences in the frequency of associations (some individuals associate often with one another, whereas others more rarely) rather than the number of associates. Frequent associations can provide experience for central females to assess the type and consistency of partner’s responses to interactions in diverse social and ecological contexts, and thereby to better assess and avoid risky situations of intra-group aggression and reduce the costs of increased vigilance in risky social contexts (Busia et al. 2019).

Contrary to our prediction, we did not find evidence for higher male versus female offspring survival for more socially integrated females. Rather, our results point to a relationship between social integration and higher female offspring survival. Although we controlled for female age by introducing reproductive years as an offset term in our models, the exclusion of five of the younger females in the multiplex analyses might have left out variation in offspring survival representing less experienced and typically less socially integrated females with a higher propensity to lose male offspring, for example, because of differences in maternal-care allocated to males during their early life-stages by younger females (Soben et al. 2023). Even so, our results suggest as much (if not higher) female offspring survival for females who are more socially integrated, and more female offspring survival for females with more stability in their integration over time. Our results regarding offspring sex suggest that social integration and its stability are more relevant to the survival of female than male offspring. As males reach sexual maturity, they develop increasingly stronger social relationships with other males (Slater et al. 2009), which might make them less dependent on their mother’s social integration compared to female offspring prior to migration.

By examining changes in social integration, we assessed whether differences in the integration process were related to differences in reproductive success as observed, for example, in spotted hyenas (Turner et al. 2021). After dispersal, spider monkey females typically undergo a progressive integration process in their new group that is reflected in low centrality/versatility scores for recent immigrants (Ramos-Fernández et al. 2009; Smith-Aguilar et al. 2019) that tends to increase as group tenure increases. Contrary to our predictions, we did not find any relationships between offspring survival or production and the change in strength or eigenvector centrality during the observation period, suggesting that variation in females’ integration process was not relevant for their reproductive success, at least to the extent captured by our metrics. However, the significant effect of residence status on offspring survival indicates that females who were already residents at the beginning of the study (and assumed to be fully integrated into the group) had higher offspring survival than females who immigrated into the group during the study. This adds to the possible effect of rearing experience on offspring survival as all the already “residents” had one or more offspring by the onset of the study. Because of high correlations between our predictors of social integration and rearing experience (e.g., measured as the number of offspring that survive to 1 year), we could not investigate their unique relationships.

While we found consistent relationships between social integration and offspring survival, it was not the case for offspring production. Female variability in social integration as captured by our indicators may not be relevant to the variability in the production of offspring, which in turn may be largely influenced by environmental factors like drought (Campos et al. 2020a) or mean temperature as observed in other ateline monkeys (Wiederholt and Post 2011).

We used two network approaches for social integration, complementing the centrality measures derived from monoplex networks with versatility measures from temporal multiplex networks. The only two differences in results observed between the approaches resulted from excluding the five females with tenures under 5 years from the multiplex analyses. It is unclear how the variability contributed by females with shorter tenure (and potentially less care-giving experience) may obscure the relationship between overall social integration and offspring survival to 1 year. As mentioned before, perhaps variation in caregiving only becomes consequential to early offspring survival after a few years of experience. Further analyses need to be conducted to clearly understand why excluding the recent immigrants makes a difference. Altogether, results from the multiplex metrics were consistent with monolayer metrics in an overall multilayer framework that provided evidence on the role of social integration for reproductive success in a female-dispersing species with a high degree of fission-fusion dynamics.

## Conclusion

Our study of the relationship between social integration and reproductive success contributes to our understanding of the benefits of social connectivity (Ostner and Schülke, 2018) with findings that add to growing evidence linking variation in social integration over time to differences in reproductive success (Turner et al. 2021). While our analyses do not establish causal relationships, they do show evidence that female spider monkeys have higher offspring survival if they are highly socially integrated and if they have more stable integration scores over time. Well-connected females may provide their offspring with better access to ecological and social resources, increasing their survival. Given the high degree of fission-fusion dynamics, the frequent changes in subgroup composition influence the patterning of social connections in a way that may be particularly relevant for information sharing about ecological and social resources. We focused on spider monkeys, but it is likely that our results are relevant more broadly to species with a high degree of fission-fusion dynamics and female dispersal.

## Supporting information

Electronic supplementary information

## Acknowledgements

We thank all field assistants (Augusto Canul, Macedonio Canul, Juan Canul and Eulogio Canul) and students who collected data during the study period. We are especially grateful to Laura Vick for sharing the management of the field site over the years. We thank the participants of the Seminar on Social Complexity for useful discussions and ideas for the development of our analyses. We also thank Secretaría de Ciencias, Humanidades, Tecnología e Innovación (SECIHTI) for providing a postdoctoral scholarship to the first and second authors under the grant CF-2019-263958 Ciencia de Frontera. Additionally, we thank the Comisión Nacional de Áreas Naturales Protegidas (CONANP), Pronatura and Secretaría de Medio Ambiente y Recursos Naturales (SEMARNAT) for permission and support to carry out the project. SSA also thanks Ruth Orozco for her invaluable support throughout the development of the study.

## Author contributions

Conceptualization: CJT, GRF, SSA; Data analyses: CJT, FA, GRF, SC, SRV, SSA, XSA; Preparation of the first draft: CTJ, SSA; Review and editing: CJT, CMS, FA, GRF, SC, SRV, SSA; Funding acquisition: CMS, FA, GRF; Supervision: GRF.

## Ethics declaration

### Ethical approval

This research adhered to Mexican law and complied with protocols approved by the Mexican environmental authority (SEMARNAT), under research permits from the Dirección General de Vida Silvestre (00910/13, 02716/14, 10405/15 and 03005/19). These permits authorized our research on a wild population of Geoffroy’s spider monkeys (an endangered species) within a protected area in Mexico for the duration of our study. The study also adhered to the ASAB/ABS Guidelines for the ethical treatment of nonhuman animals in research. Data collection was exclusively observational; therefore, none of the researchers had any direct interaction with the studied animals that could cause disturbance. The study group was continuously studied from 1997 through 2020, and thus, the researchers were a normal presence for the study animals.

## Data availability statements

Predictor datasets and R code used to fit GLMs and create plots are available as supplementary material.

