## Supplementary material for "Social integration in temporal multiplex association networks predicts offspring survival in female Geoffroy’s spider monkeys (*Ateles geoffroyi*)": Electronic supplementary information

**Electronic supplementary information for**

### Supplementary Figure S1

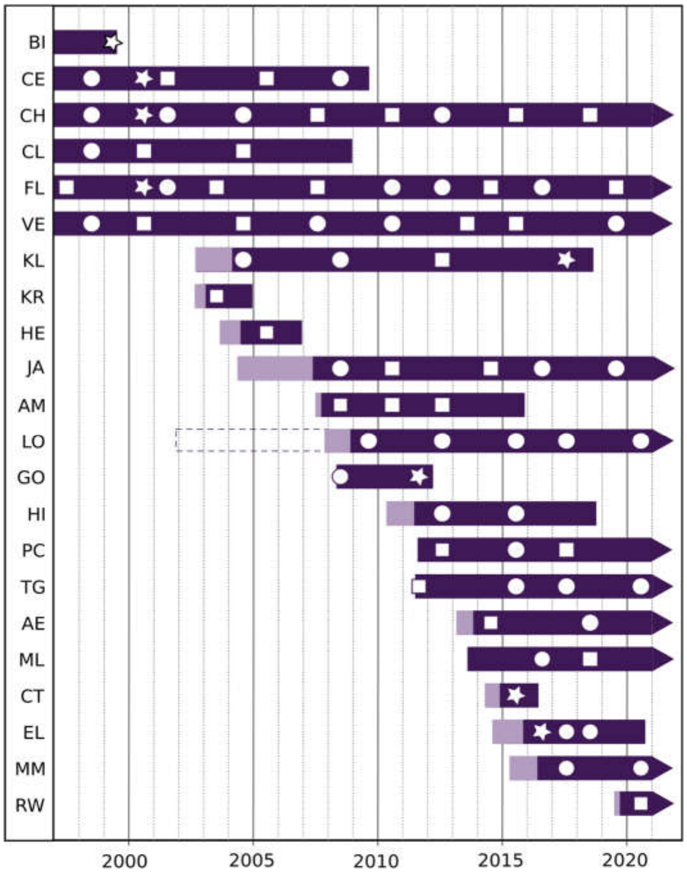


**Fig S1** Female tenure in the study group. Each bar represents one of the 22 females considered in the study, with her two-letter ID code on the left side. The left end of the bars indicates the year the female immigrated into the group or the onset of data collection for the six females already present in the group by 1997 (top rows). The right end marks the year when a female was last observed due to death or disappearance (flat edge) or the end of the study period (arrow). Bar colours distinguish between non-reproductive (lighter) and reproductive (darker) periods considered for each female. The latter were defined to begin seven months prior to the birth of the first offspring (see Table 1 for details). Inside each bar, the birth of each female's offspring is shown by a circle for female offspring, a square for male offspring or a star for offspring of unknown sex. Female LO was born in the study group and did not emigrate, reproducing in her natal group; the dashed line indicates the period of her life not considered in the study (from birth until she reached 6 years). Vertical lines mark the first day of the corresponding year

### SI. Supplementary information: Edge-weight models

Due to the difficulty to evenly sample individuals because of the high degree of fission-fusion dynamics and the patterns of immigration and disappearance, it was necessary to account for the uncertainty derived from sampling differences among monkey dyads. For this, we built Bayesian edge-weight models for each yearly network following the BiSoN framework proposed by Hart et al. (2023) for binary group-based (gambit of the group) data, defining the edge model as follows:

$$X_{ij}^{(n)} \sim Bernoulli\ (p_{ij}^{(n)})$$

$$logit\ (p_{ij}^{(n)})\ = \omega_{ij} + G_{ij}^{(n)}$$

$$G_{ij}^{(n)} \sim Normal\ (\mu_G, \sigma_G^2)$$

where $X_{ij}^{(n)}$ is the presence/absence of association between individuals *i* and *j* in the *n*-th observation of *i* and *j* (during the period in which individuals *i* and *j* coexisted in the group) over all the individuals (total= *M*) observed in a given year, resulting in an *M*x*M* association matrix (excluding pairs *i*=*j* or self-links), $p_{ij}^{(n)}$ is the probability of an association occurring between *i* and *j* in the *n*-th observation of *i* and *j*, $\omega_{ij}$ is an edge weight parameter between *i* and *j,* and $G_{ij}^{(n)}$ is a random group-level effect for the *n*-th grouping that *i* and *j* were seen in, following a normal distribution with mean $\mu_G$ and variance $\sigma_G^2$. $G_{ij}^{(n)}$ accounts for non-independence of associations occurring in the same observation (i.e. the same instantaneous scan sample). Edge-weight models were built using an integrated nested Laplace approximation (INLA) for approximate Bayesian inference implemented with the INLA package (Martino and Rue, 2009). The model included a fixed effect for dyad identity which determines edge weight estimates, and a random effect over scan sample identity. We used weakly informative priors with mean $\mu$ = 0 and variance $\sigma^2$ = 1 for both fixed and random effects. From the network, we obtained posterior distributions of edge weights for each dyad, from which we drew 1000 samples per yearly network to use in downstream analyses. Table S1 presents the number of individuals (nodes) and dyads in each yearly network.

### Supplementary Table S1

**Table S1** Size of each yearly association network in the number of nodes and dyads considered in each edge-weight model.

| Yearly network | Number of individuals (nodes) included | Number of dyads (edges) |
|---|---|---|
| 1997 | 9 | 36 |
| 1998 | 11 | 55 |
| 1999 | 10 | 45 |
| 2000 | 9 | 36 |
| 2001 | 11 | 55 |
| 2002 | 12 | 65 |
| 2003 | 13 | 78 |
| 2004 | 16 | 119 |
| 2005 | 11 | 55 |
| 2006 | 11 | 55 |
| 2007 | 15 | 105 |
| 2008 | 16 | 120 |
| 2009 | 14 | 89 |
| 2010 | 17 | 135 |
| 2011 | 16 | 120 |
| 2012 | 16 | 119 |
| 2013 | 22 | 226 |
| 2014 | 24 | 271 |
| 2015 | 26 | 324 |
| 2016 | 22 | 231 |
| 2017 | 21 | 210 |
| 2018 | 24 | 276 |
| 2019 | 24 | 267 |
| 2020 | 26 | 325 |

### Supplementary Tables S2 and S3

**Table S2** GLM results of the effect of indicators of social integration on indicators of female offspring survival, with the corresponding 95% confidence intervals (CI) pooled across 1000 Bayesian-generated social network metrics. Only predictors with at least one model with a pooled p value < 0.06 (at least marginally significant) are shown. Results for other models that were not statistically significant are presented in Table S3. Alternate rows are shaded to help identify separate models. Note that models with the Δ Strength centrality and Δ Eigenvector centrality include 2 predictors and an interaction.

| **Offspring survival variables** | ***Estimate*** | ***Std. Error*** | ***95 % CI*** | ***t value*** | ***P*** |
|---|---|---|---|---|---|
| **Number of offspring who reached 1 year** | | | | | |
| Strength versatility | 0.552 | 0.202 | (0.117, 0.987) | 2.739 | **0.017** |
| Eigenvector versatility | 0.475 | 0.121 | (0.212, 0.738) | 3.915 | **0.002** |
| $\bar{x}$ strength centrality | 0.353 | 0.205 | (-0.078, 0.784) | 1.720 | 0.102 |
| CV strength centrality | -0.003 | 0.002 | (-0.006, 0.001) | -1.759 | 0.096 |
| $\bar{x}$ eigenvector centrality | 0.323 | 0.206 | (-0.109, 0.755) | 1.569 | 0.134 |
| CV eigenvector centrality | -0.001 | 0.001 | (-0.004, 0.002) | -0.843 | 0.410 |
| Δ Strength centrality | 0.267 | 0.152 | (-0.066, 0.600) | 1.754 | 0.106 |
| Status (resident vs immigrant) | 0.161 | 0.091 | (-0.037, 0.358) | 1.772 | 0.102 |
| Δ Strength centrality* Status (resident vs immigrant) | -0.510 | 0.392 | (-1.368, 0.348) | -1.299 | 0.219 |
| Δ Eigenvector centrality | 0.210 | 0.158 | (-0.135, 0.555) | 1.329 | 0.209 |
| Status (resident vs immigrant) | 0.170 | 0.082 | (-0.010, 0.349) | 2.059 | **0.062** |
| Δ Eigenvector centrality*Status (resident vs immigrant) | -0.538 | 0.459 | (-1.540, 0.464) | -1.171 | 0.265 |
| **Number of offspring who reached 3 years** | | | | | |
| Strength versatility | 1.046 | 0.350 | (0.292, 1.800) | 2.991 | **0.010** |
| Eigenvector versatility | 0.631 | 0.257 | (0.075, 1.187) | 2.453 | **0.029** |
| $\bar{x}$ strength centrality | 0.859 | 0.309 | (0.211, 1.507) | 2.783 | **0.012** |
| CV strength centrality | -0.006 | 0.003 | (-0.012, -0.001) | -2.423 | **0.027** |
| $\bar{x}$ eigenvector centrality | 0.803 | 0.314 | (0.144, 1.463) | 2.559 | **0.020** |
| CV eigenvector centrality | -0.005 | 0.002 | (-0.009, 0.0001) | -2.163 | **0.045** |
| Δ Strength centrality | 0.312 | 0.320 | (-0.384, 1.009) | 0.976 | 0.348 |
| Status (resident vs immigrant) | 0.278 | 0.188 | (-0.131, 0.688) | 1.482 | 0.165 |

| | | | | | |
|---|---|---|---|---|---|
| Δ Strength centrality*Status (resident vs immigrant) | -0.695 | 0.798 | (-2.441, 1.052) | -0.871 | 0.402 |
| Δ Eigenvector centrality | 0.306 | 0.318 | (-0.386, 0.998) | 0.961 | 0.355 |
| Status (resident vs immigrant) | 0.293 | 0.165 | (-0.067, 0.653) | 1.769 | 0.102 |
| Δ Eigenvector centrality*Status (resident vs immigrant) | -1.061 | 0.910 | (-3.057, 0.935) | -1.166 | 0.268 |
| **Number of offspring who reached 5 years** | | | | | |
| Strength versatility | 2.250 | 0.568 | (1.025, 3.474) | 3.962 | **0.002** |
| Eigenvector versatility | 1.351 | 0.462 | (0.350, 2.352) | 2.922 | **0.012** |
| $\overline{x}$ strength centrality | 2.094 | 0.425 | (1.203, 2.985) | 4.932 | **0.0001** |
| CV strength centrality | -0.013 | 0.005 | (-0.023, -0.003) | -2.758 | **0.013** |
| $\overline{x}$ eigenvector centrality | 2.065 | 0.436 | (1.150, 2.980) | 4.738 | **0.0002** |
| CV eigenvector centrality | -0.010 | 0.004 | (-0.019, -0.002) | -2.534 | **0.021** |
| Δ Strength centrality | 0.492 | 0.526 | (-0.654, 1.637) | 0.935 | 0.368 |
| Status (resident vs immigrant) | 0.962 | 0.273 | (0.369, 1.556) | 3.530 | **0.004** |
| Δ Strength centrality*Status (resident vs immigrant) | -0.674 | 1.023 | (-2.901, 1.554) | -0.659 | 0.523 |
| Δ Eigenvector centrality | 0.588 | 0.509 | (-0.521, 1.696) | 1.154 | 0.271 |
| Status (resident vs immigrant) | 0.967 | 0.249 | (0.426, 1.509) | 3.882 | **0.002** |
| Δ Eigenvector centrality*Status (resident vs immigrant) | -1.100 | 1.129 | (-3.561, 1.362) | -0.974 | 0.349 |
| **Number of female offspring who reached 5 years** | | | | | |
| Strength versatility | 3.308 | 0.744 | (1.701, 4.915) | 4.445 | **0.001** |
| Eigenvector versatility | 2.030 | 0.638 | (0.647, 3.412) | 3.183 | **0.007** |
| $\overline{x}$ strength centrality | 2.752 | 0.672 | (1.341, 4.163) | 4.094 | **0.001** |
| CV strength centrality | -0.022 | 0.008 | (-0.039, -0.005) | -2.788 | **0.012** |
| $\overline{x}$ eigenvector centrality | 2.654 | 0.706 | (1.171, 4.136) | 3.758 | **0.001** |
| CV eigenvector centrality | -0.019 | 0.007 | (-0.032, -0.005) | -2.826 | **0.011** |
| Δ Strength centrality | 0.446 | 0.880 | (-0.654, 1.637) | 0.506 | 0.622 |
| Status (resident vs immigrant) | 1.242 | 0.433 | (0.369, 1.556) | 2.865 | **0.014** |
| Δ Strength centrality*Status (resident vs immigrant) | -0.042 | 1.627 | (-2.901, 1.554) | -0.026 | 0.980 |
| Δ Eigenvector centrality | 0.500 | 0.885 | (-0.521, 1.696) | 0.565 | 0.582 |
| Status (resident vs immigrant) | 1.189 | 0.414 | (0.426, 1.509) | 2.873 | **0.014** |
| Δ Eigenvector centrality*Status (resident vs immigrant) | -0.869 | 1.816 | (-3.561, 1.362) | -0.479 | 0.641 |
| **Number of male offspring who reached 5 years** | | | | | |
| Strength versatility | 1.152 | 0.943 | (-0.880, 3.184) | 1.222 | 0.243 |
| Eigenvector versatility | 0.683 | 0.667 | (-0.756, 2.122) | 1.023 | 0.325 |
| $\overline{x}$ strength centrality | 1.458 | 0.686 | (0.019, 2.897) | 2.127 | **0.047** |
| CV strength centrality | -0.006 | 0.006 | (-0.018, 0.006) | -1.062 | 0.303 |

| | | | | | |
|---|---|---|---|---|---|
| $\bar{x}$ eigenvector centrality | 1.496 | 0.698 | (0.031, 2.962) | 2.143 | **0.046** |
| CV eigenvector centrality | -0.004 | 0.005 | (-0.014, 0.006) | -0.903 | 0.379 |
| Δ Strength centrality | 0.526 | 0.832 | (-1.286, 2.338) | 0.632 | 0.539 |
| Status (resident vs immigrant) | 0.661 | 0.460 | (-0.340, 1.663) | 1.437 | 0.176 |
| Δ Strength centrality*Status (resident vs immigrant) | -1.400 | 1.755 | (-5.232, 2.432) | -0.798 | 0.441 |
| Δ Eigenvector centrality | 0.655 | 0.824 | (-1.138, 2.448) | 0.795 | 0.442 |
| Status (resident vs immigrant) | 0.741 | 0.423 | (-0.178, 1.660) | 1.753 | 0.105 |
| Δ Eigenvector centrality*Status (resident vs immigrant) | -1.344 | 2.017 | (-5.746, 3.058) | -0.667 | 0.518 |

Below each indicator of female reproductive success (see Table 1 for definitions), reported in bold, we show the result for each social integration predictor (see Table 2 for definitions) from each GLM.

**Table S3** Non-significant GLM results of the effect of each indicator of social integration on indicators of offspring production with the corresponding 95% CI pooled across 1000 Bayesian-generated social network metrics. Alternate rows are shaded to help identify separate models. Note that models with the Δ Strength centrality and Δ Eigenvector centrality include 2 predictors and an interaction.

| **Offspring production variables** | ***Estimate*** | ***Std. Error*** | ***95 % CI*** | ***t value*** | ***P*** |
|---|---|---|---|---|---|
| **Number of offspring** | | | | | |
| Strength versatility | 0.171 | 0.284 | (-0.442, 0.783) | 0.601 | 0.558 |
| Eigenvector versatility | 0.257 | 0.188 | (-0.149, 0.663) | 1.364 | 0.195 |
| $\bar{x}$ strength centrality | 0.048 | 0.246 | (-0.469, 0.564) | 0.193 | 0.849 |
| CV strength centrality | 0.000 | 0.002 | (-0.004, 0.004) | -0.057 | 0.955 |
| $\bar{x}$ eigenvector centrality | 0.026 | 0.244 | (-0.485, 0.537) | 0.106 | 0.917 |
| CV eigenvector centrality | -0.001 | 0.002 | (-0.004, 0.002) | -0.449 | 0.659 |
| Δ Strength centrality | 0.260 | 0.198 | (-0.171, 0.692) | 1.316 | 0.213 |
| Status (resident vs immigrant) | 0.046 | 0.121 | (-0.218, 0.310) | 0.379 | 0.711 |
| Δ Strength centrality*Status (resident vs immigrant) | -0.340 | 0.548 | (-1.539, 0.858) | -0.620 | 0.547 |
| Δ Eigenvector centrality | 0.231 | 0.202 | (-0.210, 0.671) | 1.140 | 0.276 |
| Status (resident vs immigrant) | 0.042 | 0.109 | (-0.194, 0.278) | 0.387 | 0.706 |
| Δ Eigenvector centrality*Status (resident vs immigrant) | -0.433 | 0.631 | (-1.811, 0.945) | -0.686 | 0.506 |
| **Number of female offspring** | | | | | |
| Strength versatility | 0.513 | 0.736 | (-1.073, 2.100) | 0.698 | 0.497 |
| Eigenvector versatility | 0.481 | 0.505 | (-0.608, 1.569) | 0.952 | 0.358 |

| | | | | | |
|---|---|---|---|---|---|
| $\bar{x}$ strength centrality | 0.251 | 0.525 | (-0.851, 1.353) | 0.478 | 0.638 |
| CV strength centrality | 0.001 | 0.004 | (-0.007, 0.009) | 0.243 | 0.811 |
| $\bar{x}$ eigenvector centrality | 0.128 | 0.518 | (-0.959, 1.216) | 0.248 | 0.807 |
| CV eigenvector centrality | 0.001 | 0.003 | (-0.005, 0.007) | 0.281 | 0.782 |
| Δ Strength centrality | 0.383 | 0.523 | (-0.755, 1.521) | 0.733 | 0.478 |
| Status (resident vs immigrant) | 0.036 | 0.330 | (-0.682, 0.754) | 0.108 | 0.915 |
| Δ Strength centrality*Status (resident vs immigrant) | 0.182 | 1.586 | (-3.292, 3.656) | 0.115 | 0.911 |
| Δ Eigenvector centrality | 0.312 | 0.537 | (-0.855, 1.480) | 0.582 | 0.571 |
| Status (resident vs immigrant) | -0.034 | 0.298 | (-0.681, 0.614) | -0.113 | 0.912 |
| Δ Eigenvector centrality*Status (resident vs immigrant) | -0.358 | 1.799 | (-4.300, 3.585) | -0.199 | 0.846 |
| **Number of male offspring** | | | | | |
| Strength versatility | -0.107 | 0.787 | (-1.804, 1.591) | -0.136 | 0.894 |
| Eigenvector versatility | 0.099 | 0.541 | (-1.066, 1.265) | 0.183 | 0.857 |
| $\bar{x}$ strength centrality | -0.021 | 0.611 | (-1.303, 1.261) | -0.034 | 0.973 |
| CV strength centrality | -0.006 | 0.005 | (-0.016, 0.005) | -1.073 | 0.298 |
| $\bar{x}$ eigenvector centrality | 0.013 | 0.608 | (-1.263, 1.289) | 0.021 | 0.983 |
| CV eigenvector centrality | -0.003 | 0.004 | (-0.011, 0.006) | -0.657 | 0.520 |
| Δ Strength centrality | 0.125 | 0.647 | (-1.283, 1.533) | 0.193 | 0.850 |
| Status (resident vs immigrant) | 0.084 | 0.382 | (-0.747, 0.915) | 0.221 | 0.829 |
| Δ Strength centrality*Status (resident vs immigrant) | -1.093 | 1.606 | (-4.593, 2.407) | -0.681 | 0.509 |
| Δ Eigenvector centrality | 0.126 | 0.655 | (-1.298, 1.550) | 0.192 | 0.851 |
| Status (resident vs immigrant) | 0.173 | 0.336 | (-0.558, 0.904) | 0.514 | 0.617 |
| Δ Eigenvector centrality*Status (resident vs immigrant) | -0.809 | 1.867 | (-4.882, 3.264) | -0.433 | 0.672 |

Below each indicator of female reproductive success (see Table 1 for definitions), reported in bold, we show the result for each social integration predictor (see Table 2 for definitions) from each GLM.

**Supplementary Figure S2**

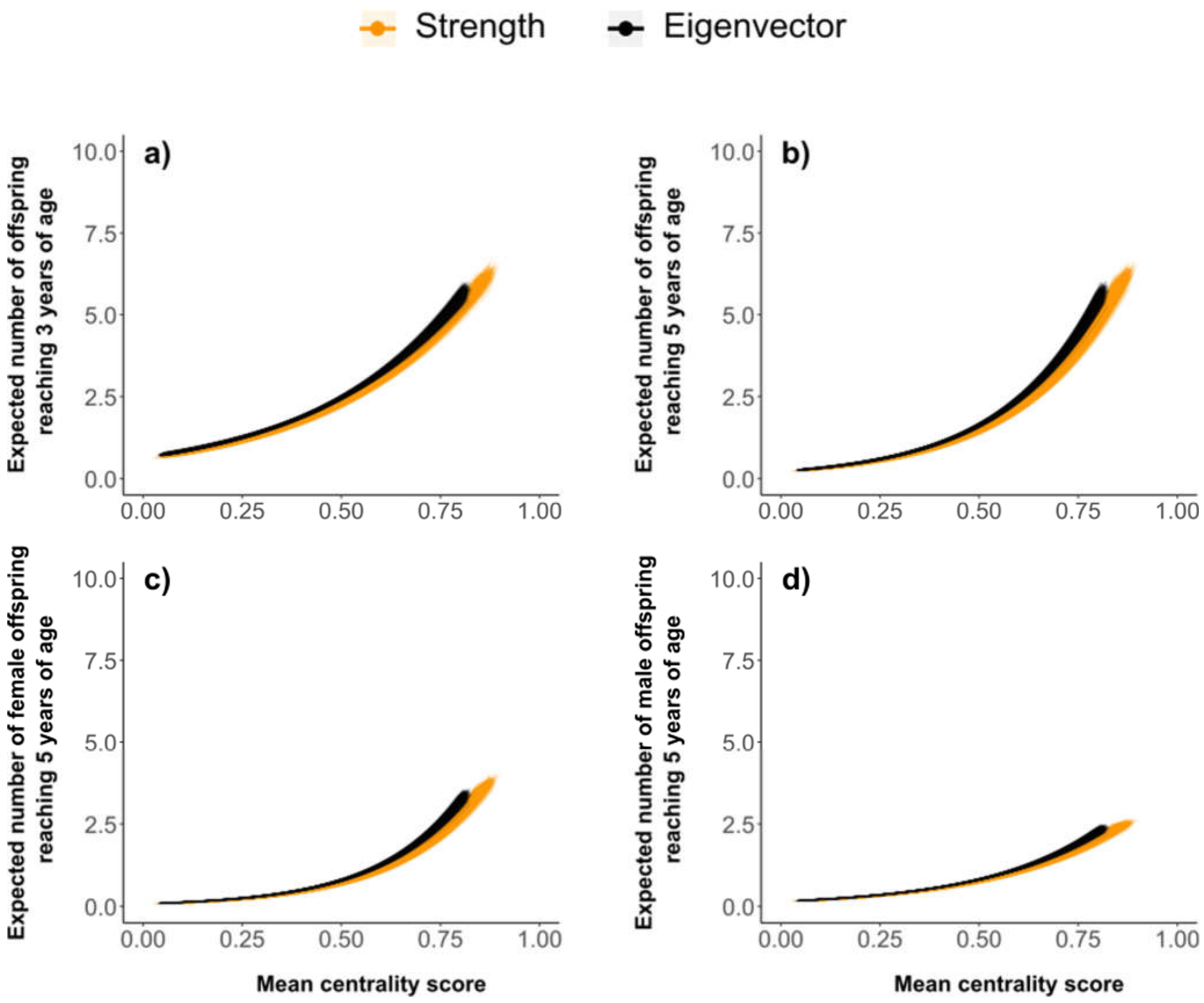


**Fig S2** GLM quasi-poisson trend lines (curves) for mean strength (orange) and eigenvector centrality (black) showing their relationship with the expected number of offspring to survive to 3 (a) and 5 years (b), and the expected number of female (c) and male (d) offspring to survive to 5 years. Each trend line represents one of the 1,000 GLM models
